# Identification of multiple TAR DNA Binding Protein retropseudogene lineages during the evolution of primates

**DOI:** 10.1101/2021.05.24.445516

**Authors:** Juan C. Opazo, Kattina Zavala, Luis Vargas-Chacoff, Francisco J. Morera, Gonzalo A. Mardones

**Author notes:** **Correspondence:** Juan C. Opazo, Gonzalo A. Mardones.

## Abstract

The TAR DNA Binding Protein (TARDBP) gene has gained attention in biomedicine after the discovery of several pathogenic mutations. The lack of knowledge about its evolutionary history contrasts with a large number of studies in the biomedical area. This study aimed to investigate the retrotransposition evolutionary dynamics associated with this gene in primates. We identified retropseudogenes that originated in the ancestors of anthropoids, catarrhines, and lemuriformes, i.e. the strepsirrhine clade that inhabit Madagascar. We also found species-specific retropseudogenes in the philippine tarsier, Bolivian squirrel monkey, capuchin monkey and vervet. Although retropseudogenes are not able to produce a functional protein, we can not rule out that they may represent genetic material upon which evolution acts on, especially with regulatory functions.

## Introduction

Today, the availability of whole-genome sequences allows performing more detailed research on the evolution of different genetic elements. An important source of evolutionary innovation are the events of RNA retrotranscription and its insertion into the genome (Kaessmann et al. 2009; Casola & Betrán 2017). These events depend on the enzymatic machinery encoded by retrotransposable elements, and they produce an intronless gene duplicate that could produce a protein similar to the parental gene (Zhang 2003). However, most retrotranscribed sequences are inserted at a random position in the genome, lacking all necessary transcription elements and becoming a pseudogene, a phenomenon called “dead on arrival” (Zhang 2003). However, retroseudogenes are not necessarily a dead end. There are cases in which they encode small interfering RNAs with regulatory functions (Tam et al. 2008; Watanabe et al. 2008). Thus, identifying the presence of retrocopies/retroseudogenes is an important piece of information to have a complete picture of the evolution of any particular gene.

The TAR DNA Binding Protein (TARDBP) gene, which encodes the Transactive response DNA-binding protein 43 kDa (TDP-43), has gained considerable attention after the initial discovery that its mutations can cause familial amyotrophic lateral sclerosis (ALS) and frontotemporal dementia (FTD), two major forms of neurodegenerative disorders (Sreedharan et al. 2008; Kabashi et al. 2008). Up to date, more than 50 pathogenic missense mutations have been characterized (Sreedharan et al. 2008; Kabashi et al. 2008). TDP-43 is an RNA-binding protein with a variety of RNA metabolism functions, including transcription, mRNA transport and stabilization, miRNA biogenesis, long noncoding RNA processing, and translation (Hanson et al. 2012). More recent findings indicate that TDP-43 participates in the pathogenesis of other neurodegenerative disorders of several other proteinopathies, such as Parkinson’s disease and Alzheimer’s disease, which are conditions characterized by toxic protein aggregation (Klim et al. 2021). In human cells, under physiological conditions, TDP-43 mainly localizes in the nucleus, but in neurons and glial cells of ALS and FTD patients it shuttles and accumulates in the cytoplasm where eventually aggregates and contribute to the onset and progression of these diseases (Neumann et al. 2006; Arai et al. 2006; Robberecht & Philips 2013; Heyburn & Moussa 2017; Pinarbasi et al. 2018). The TARDBP gene is conserved in species that share a common ancestor deep in time (Wang et al. 2004), suggesting that this gene carries out essential functions. This gene underwent an event of positive selection in the ancestor of mammals (Zhao et al. 2020), suggesting functional adaptations for the group. More recently in evolutionary time, it has been shown that during the evolution of humans, genes related to diseases like Alzheimer’s also underwent positive selection (Vamathevan et al. 2008). Although events of positive selection are seen as conferring selective advantage, as a by-product, they can also have adverse effects (Holt et al. 1996). In this regard, it is proposed that human susceptibility to neurodegenerative disorders could be a consequence of improving our cognitive function (Gearing et al. 1994; Keller & Miller 2006).

The aim of this study is to investigate the retrotransposition dynamics associated with the TARDBP gene in primates. According to our phylogenetic and synteny analyses, we identified retropseudogenes that originated at different times during the evolution of primates. TARDBP retropseudogenes originated in the anthropoid ancestor, between 67 and 43.2 million years ago, in the ancestor of catarrhines, between 43.2 and 29.44 million years ago, and in the ancestor of lemuriformes, i.e. the strepsirrhine clade that inhabit madagascar, between 59.3 and 55 million years ago. We also found species-specific retropseudogenes in the philippine tarsier (*Carlito syrichta*), Bolivian squirrel monkey (*Saimiri boliviensis*), capuchin monkey (*Cebus capucinus imitator*) and vervet (*Chlorocebus sabaeus*). Although annotated sequences are not putatively functional, given the presence of indels and premature stop codons, we can not rule out that these retrotransposed sequences may represent material on which evolution acts, especially to regulate the expression of their parental gene.

## Results

### Multiple retropseudogenes lineages characterize the evolution of the TARDBP gene in primates

According to our phylogenetic and synteny analyses, we identified retropseudogenes of the TARDBP gene that originated at different times during the evolution of primates. We identified retropseudogenes originated in the ancestor of anthropoids, between 67 and 43.2 million years ago, in the ancestor of catarrhines, between 43.2 and 29.44 million years ago, and in the ancestor of lemuriformes, i.e. the strepsirrhine clade that inhabit madagascar, between 59.3 and 55 million years ago (Fig. 1). More recently in evolutionary time, we found species-specific retropseudogenes in the philippine tarsier (*Carlito syrichta*), Bolivian squirrel monkey (*Saimiri boliviensis*), capuchin monkey (*Cebus capucinus imitator*) and vervet (*Chlorocebus sabaeus*). All of them did not have intron sequences and were identified on a different autosome in comparison to the chromosomal location of the functional copy (Fig. 1). Our gene tree did not significantly deviate from the most updated phylogenetic hypotheses for the main group of primates (Pozzi et al. 2014; Finstermeier et al. 2013; Perelman et al. 2011), suggesting that the functional copy of the TARDBP gene was present as a simple copy gene in the ancestor of the group (Fig. 1).

**Figure 1.**
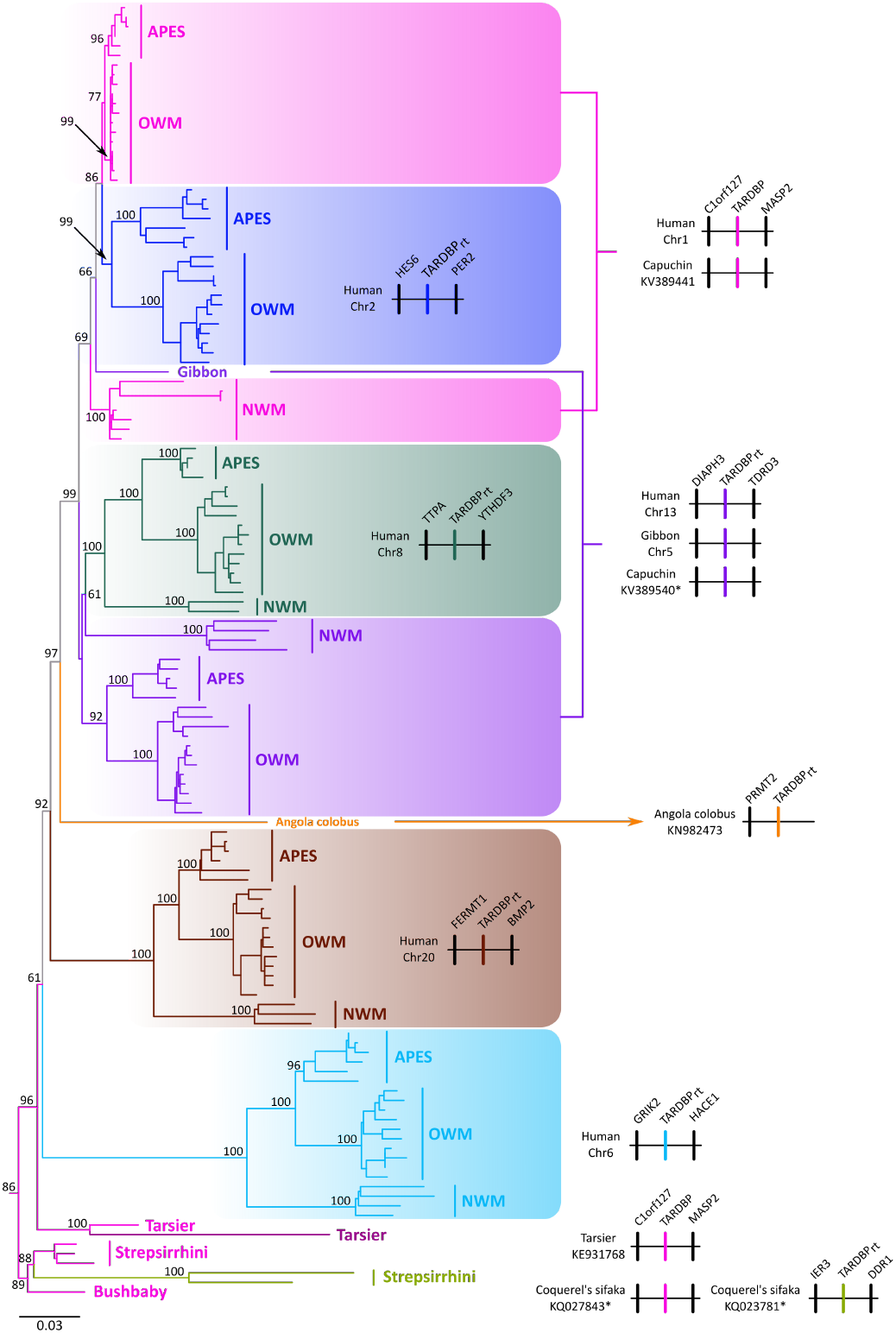
Maximum likelihood tree showing sister group relationships between the functional copy of TARDBP and primate retropseudogenes. Numbers on the nodes correspond to support values from the ultrafast bootstrap routine. TARDBP sequences from the African elephant (*Loxodonta africana*), blue whale (*Balaenoptera musculus*) and red fox (*Vulpes vulpes*) were used as outgroups (not shown). The scale denotes substitutions per site and colors represent lineages. The pink lineage represents the TARDBP functional copy. Synteny information is provided for each lineage at the right side of the figure.

We recovered three highly supported monophyletic groups containing representative species of all major groups of anthropoids i.e. apes, Old World monkeys and New World monkeys (light blue, brown and green lineages, Fig. 1), indicating that these retropseudogenes originated in the ancestor of the group, between 67 and 43.2 millions of years ago, and were maintained in representative species of all descendant primate groups. The retropseudogene lineage depicted with the purple shading (Fig. 1), although it was not recovered monophyletic, our synteny analyses suggest that it indeed belongs to a single lineage (Fig. 1). Representative species of the three purple clades possess the same flanking genes, DIAPH3 at the 5’ side and TDRD3 at the 3’ side of the retropseudogene, strongly suggesting that the lack of monophyly could be attributed to a phylogenetic artifact (Fig. 1). The small number of changes, as illustrated by the short branches that define the sister group relationships of the main clades, could be the main cause (Fig. 1). We also found a retropseudogene lineage that according to our phylogenetic tree originated in the ancestor of catarrhine primates, the group that includes apes and Old World monkeys (blue lineage, Fig. 1), between 43.2 and 29.44 million years ago. In this case, we recovered a clade containing the functional copy of the TARDBP gene in catarrhines (upper pink lineage, Fig. 1), sister to a group containing a retropseudogene in the same primate group (blue lineage, Fig. 1). The clade containing TARDBP functional sequences from New World monkeys was recovered sister to the above mentioned clade (Fig. 1). In this clade in addition to the functional TARDBP copy, we found New World monkey specific retropseudogenes for which the evolutionary history is difficult to resolve given the shortness of the branches (Fig. 1). We identified three retropseudogenes, two in the capuchin monkey (*Cebus capucinus imitator*) and one in the Bolivian squirrel monkey (*Saimiri boliviensis*). Finally, we recovered a sequence from the Angola colobus (*Colobus angolensis*)(*yellow* branch, Fig. 1), which was recovered sister to a clade containing the TARDBP functional copy (pink lineage, Fig. 1) and three retropseudogenes lineages (blue, purple and green clades, Fig. 1). The phylogenetic position of this branch in our gene tree suggests that it represents a retropseudogene originated in the anthropoid ancestor, but only conserved in this species. In support of this claim, the single flanking gene (PRMT2) found in the genomic piece containing the TARDBP retropseudogene in the Angola colobus is not shared with any other gene lineage described in this study (Fig. 1). In all cases, the identified retropseudogenes during the evolutionary history of anthropoid primates have premature stop codons and indels (supplementary figures 1 to 5).

We also identified retropseudogenes in tarsiers and strepsirrhines (Fig. 1). We found a single retropseudogene in the Philippine tarsier (*Carlito syrichta*), which shows the hallmark of a sequence free from selective constraints, i.e., a long branch as a signal of an accelerated rate of evolution in comparison to the functional copy (Fig. 1). In the strepsirrhine clade we identified a highly supported lineage containing the TARDBP functional copy in three species, greater bamboo lemur (*Prolemur simus*), coquerel’s sifaka (*Propithecus coquereli*) and the mouse lemur (*Microcebus murinus*), which in turn was recovered sister to an also highly supported clade containing retropseudogenes in the greater bamboo lemur (*Prolemur simus*) and coquerel’s sifaka (*Propithecus coquereli*) (Fig.1). This tree topology suggests that this retrocopy originated in the ancestor of lemuriformes, i.e. the strepsirrhine clade that inhabits Madagascar, between 59.3 and 55 million years ago, and it has been maintained in the genome of descendant species. Finally, the functional copy of the bushbaby (*Otolemur garnettii*) was recovered sister to the lemuriformes clade. Similar to the case of anthropoids, all retropseudogenes identified in tarsiers and strepsirrhines have premature stop codons and indels (supplementary figures 6 and 7).

## Discussion

In this study we revealed that the evolutionary history of TARDBP, an RNA-binding protein involved in several human neurodegenerative disorders (Sreedharan et al. 2008; Kabashi et al. 2008), is characterized by the presence of retropseudogenes that originated at different ages during the evolutionary history of primates. An important fraction of the retropseudogenes originated in the anthropoid ancestor, between 67 and 43.2 million years ago, and has remained in the genome of the species (Fig. 2). This phenomenon fits the expectation of a peak of retrocopy formation around 40 million years ago, which coincides with an increased activity of L1 retroelements that produced an explosion in SINE/Alu repeat amplification (Ohshima et al. 2003; Marques et al. 2005). Interestingly, this period of time represents a key moment during the evolutionary history of primates, the radiation of the anthropoid lineage, where significant morphological and physiological traits arouse (Kay et al. 1997).

**Figure 2.**
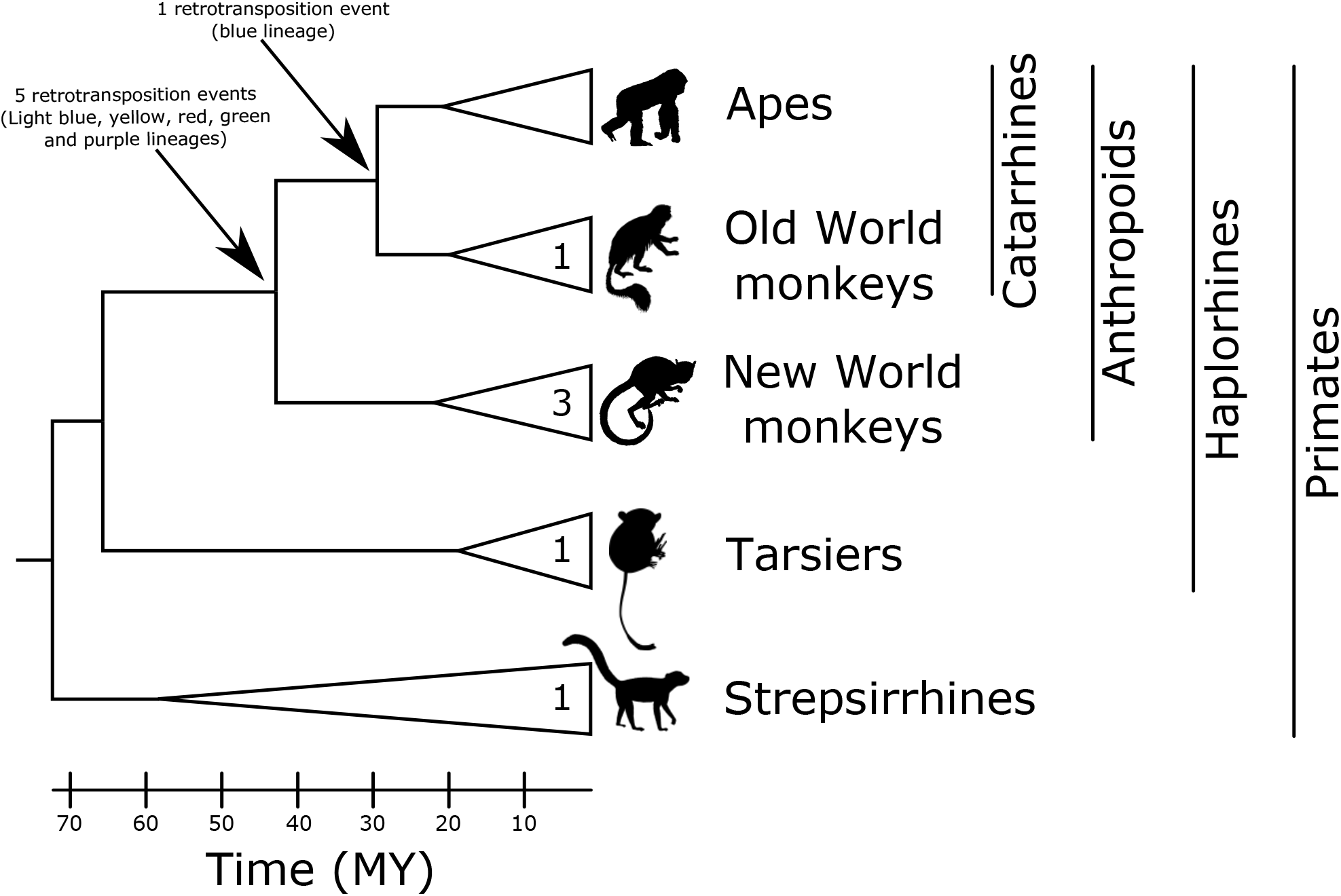
Time calibrated primate phylogeny showing the origin of the different retropseudogene lineages depicted in figure 1. Numbers on the triangles correspond to retropseudogenes originated within a particular group of primates. Silhouette images were obtained from PhyloPic (http://phylopic.org/). Divergence times were obtained from timetree (http://www.timetree.org/)(Kumar et al. 2017).

Although we did not find retrocopies, i.e. intronless copies with the potential to produce a similar protein compared to the parental gene, it has been suggested that retropseudogenes could encode small interfering RNAs that can regulate the expression of their parental genes (Tam et al. 2008; Watanabe et al. 2008). Thus, this period of Vesuvian mode of evolution could be seen as a source of evolutionary novelty that fueled the origin of the phenotypes that define the anthropoid lineage (Long et al. 2003; Kaessmann et al. 2009; Casola & Betrán 2017). Other retropseudogenes originated in the catarrhine ancestor, between 43.2 and 29.44 million years ago, and in other primate groups (Fig. 2).

In agreement with the literature, and given the nature of the process originating retrocopies, all of them seem to be non-functional as canonical TARDBP (Zhang 2003), which can be verified by the presence of indels and premature stop codons (supplementary figures 1 and 7). The identification of several retropseudogenes for the TARDBP gene in primates appears to be not a surprise as this gene comply all the requisites to be a gene with multiple retropseudogenes (Zhang & Gerstein 2004; McDonell & Drouin 2012; Gonçalves et al. 2000), i.e., short transcripts (61 to 414 amino acids)(Yates et al. 2020), widely and highly expressed (Uhlén et al. 2015), low GC-content (47%, average among 23 primate species) and highly conserved (3.37%, maximal divergence among primates). Furthermore, in agreement with slow rate of length abridgment, the identified retropseudogenes possess a length (mean 1128 bp, median 1193 bp) similar to the functional TARDBP gene (1245 bp).

Among apes, the number of TARDBP retrotransposition events appear to be higher in comparison to the average number of retrocopies per parental gene in their genomes (Navarro & Galante 2015). On average, ape genomes possess 2.91 retrocopies per parental gene (Navarro & Galante 2015), however in our study we identified five TARDBP retropseudogenes in each examined ape species. Coincident with previous evidence, we also found a higher number of retropseudogenes in New World monkeys (Navarro & Galante 2015). Although it is not clear the reason why this primate group possesses more retrocopies in comparison to catarrhines, it is suggested that a lineage specific expansion of L1PA1 and L1P3 subelements could be linked to the observed pattern (Navarro & Galante 2015; Casola & Betrán 2017).

In conclusion, in this work, we demonstrate that the TARDBP gene in primates has an evolutionary history characterized by the presence of multiple retropseudogene lineages. In the ancestor of anthropoids occurred a burst of activity, which led to intronless sequences that can not give rise to functional proteins. However, we can not discard that these DNA retrotransposed sequences could represent the raw genetic material for the evolution to act on, especially to regulate the expression of their parental gene, as it has been described for other genes.

## Material and Methods

### DNA sequences

#### DNA sequences and phylogenetic analyses

We performed searches for TAR DNA Binding Protein (TARDBP) genes in primate genomes in Ensembl v.102 (Yates et al. 2020). We retrieved primate orthologs, using the human (*Homo sapiens*) entry, based on the ortholog prediction function of Ensembl v.102 (Yates et al. 2020). We identified TARDBP retropseudogenes in primate species by performing BLASTN searches (Altschul et al. 1990), against the whole genome sequence in Ensembl v.102 (Yates et al. 2020) using default settings. In each case the query sequence (TARDBP) was from the same species of the genome in which retropseudogenes were looking for. In our searches, a retropseudogene was recognized as a sequence containing all exons together and found in a different chromosome in comparison to the functional copy. Genomic fragments containing retropseudogenes were extracted and manually annotated by comparing the coding sequence of the same species using the program Blast2seq v2.5 (Tatusova & Madden 1999) with default parameters. Accession numbers and details about the taxonomic sampling are available in Supplementary Table S1.

Nucleotide sequences were aligned using MAFFT v.7 (Katoh & Standley 2013), allowing the program to choose the alignment strategy (FFT-NS-i). We used the proposed model tool of IQ-Tree v.1.6.12 (Kalyaanamoorthy et al. 2017) to select the best-fitting model of nucleotide substitution, which selected GTR+F+R3. We used the maximum likelihood method to obtain the best tree using the program IQ-Tree v1.6.12 (Trifinopoulos et al. 2016). We assessed support for the nodes using three strategies: a Bayesian-like transformation of aLRT (aBayes test) (Anisimova et al. 2011), SH-like approximate likelihood ratio test (SH-aLRT) (Guindon et al. 2010) and the ultrafast bootstrap approximation (Hoang et al. 2018). We carried out 20 independent runs to explore the tree space, and the tree with the highest likelihood score was chosen. TARDBP sequences from the African elephant (*Loxodonta africana*), blue whale (*Balaenoptera musculus*) and red fox (*Vulpes vulpes*) were used as outgroups.

#### Assessment of conserved synteny

We examined genes found upstream and downstream of functional copies and retropseudogenes. We used the estimates of orthology and paralogy derived from the Ensembl Compara database (Herrero et al. 2016); these estimates are obtained from a pipeline that considers both synteny and phylogeny to generate orthology mappings. These predictions were visualized using the program Genomicus v100.01 (Nguyen et al. 2018). Our assessments were performed in representative species for each lineage.

## Acknowledgements

This work was supported by Fondo Nacional de Desarrollo Científico y Tecnológico from Chile (FONDECYT 1210471) and Millennium Nucleus of Ion Channels Associated Diseases (MiNICAD), Iniciativa Científica Milenio, Ministry of Economy, Development and Tourism from Chile to JCO, Fondo Nacional de Desarrollo Científico y Tecnológico from Chile (FONDECYT 1180957) to FJM and LVC and Fondo Nacional de Desarrollo Científico y Tecnológico from Chile (FONDECYT 1211481) to GAM.

## Author contributions

JCO and GAM designed the study. KZ, JCO collected and analyzed data. JCO and GAM wrote the manuscript. LV-C, FJM reviewed and edited the manuscript. All authors contributed to the article and approved the submitted version.

## Competing Interests

The authors declare no competing of interest

---

**Supplementary figure 1.**
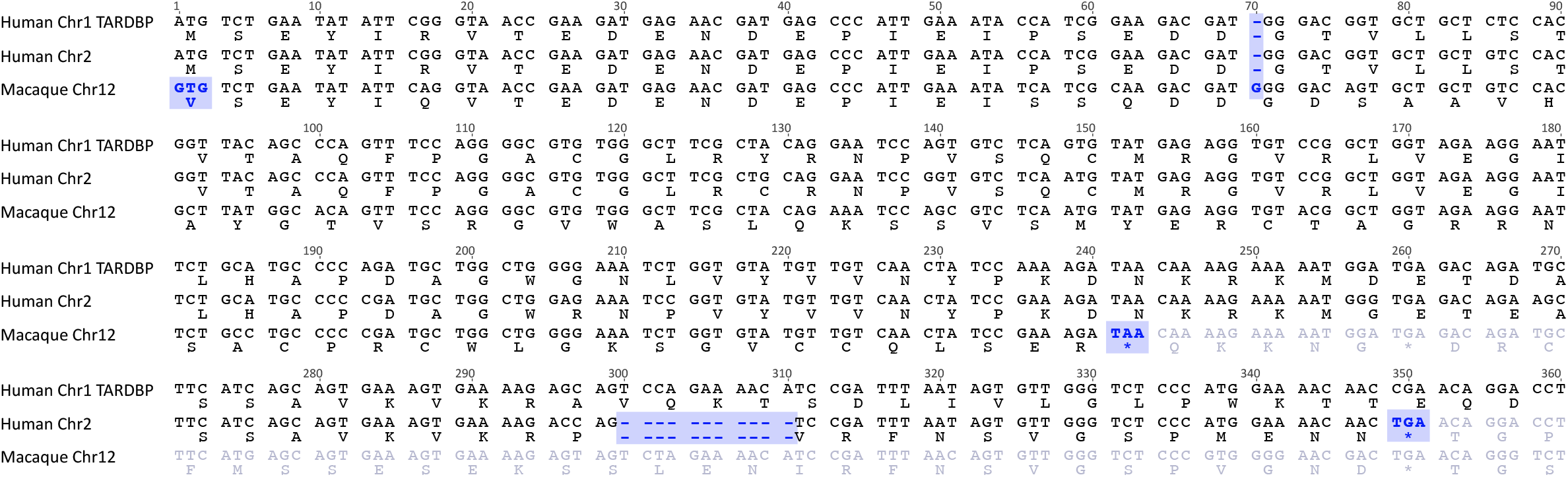
Nucleotide alignment of the TARDBP functional copy of humans (*Homo sapiens*) and TARDBP retrocopies in representative species of primates in which the retrocopy was identified corresponding to the blue lineage on figure 1. The shading highlights the mutations that make the retrocopies non-functional.

**Supplementary figure 2.**
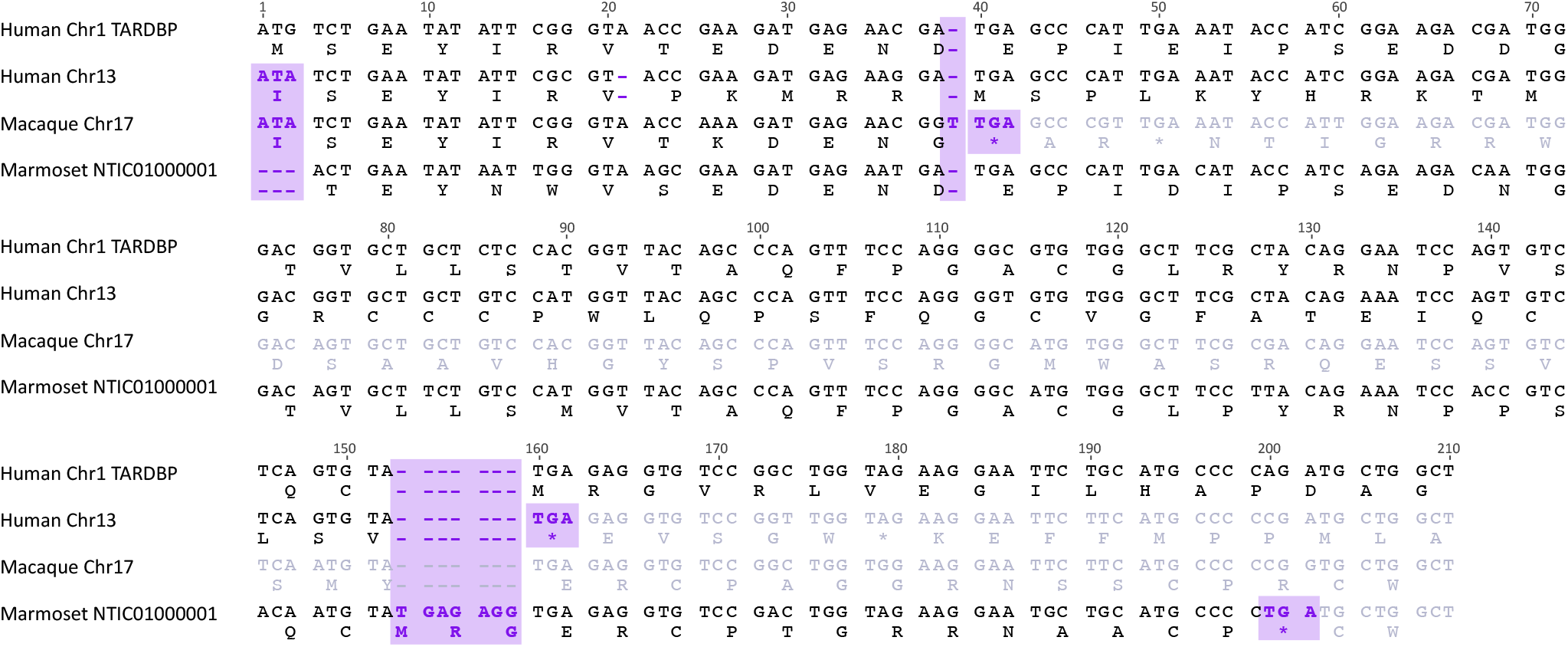
Nucleotide alignment of the TARDBP functional copy of humans (*Homo sapiens*) and TARDBP retrocopies in representative species of primates in which the retrocopy was identified corresponding to the purple lineage on figure 1. The shading highlights the mutations that make the retrocopies non-functional.

**Supplementary figure 3.**
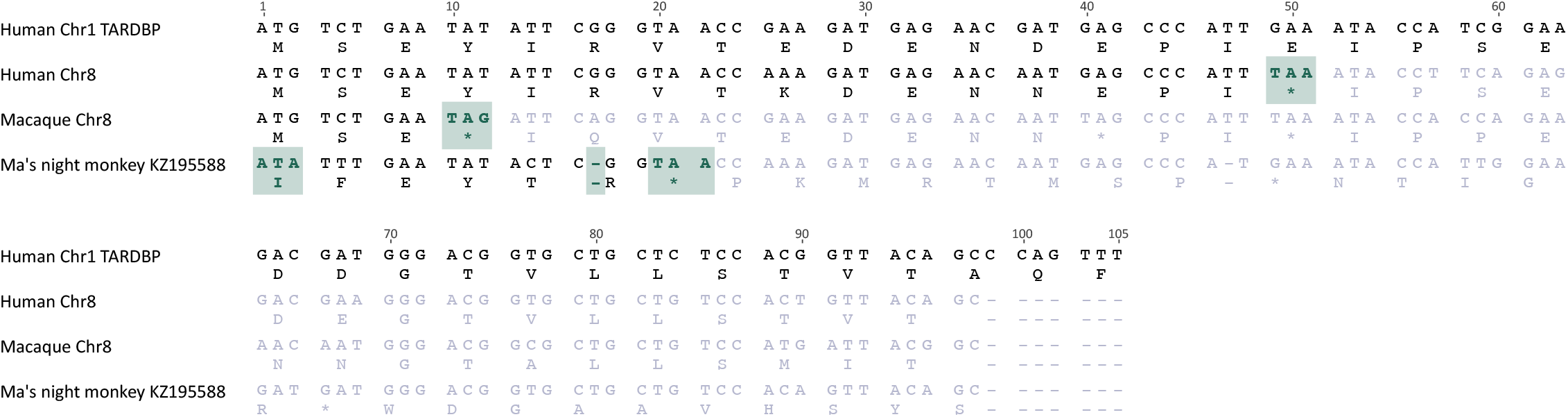
Nucleotide alignment of the TARDBP functional copy of humans (*Homo sapiens*) and TARDBP retrocopies in representative species of primates in which the retrocopy was identified corresponding to the green lineage on figure 1. The shading highlights the mutations that make the retrocopies non-functional.

**Supplementary figure 4.**
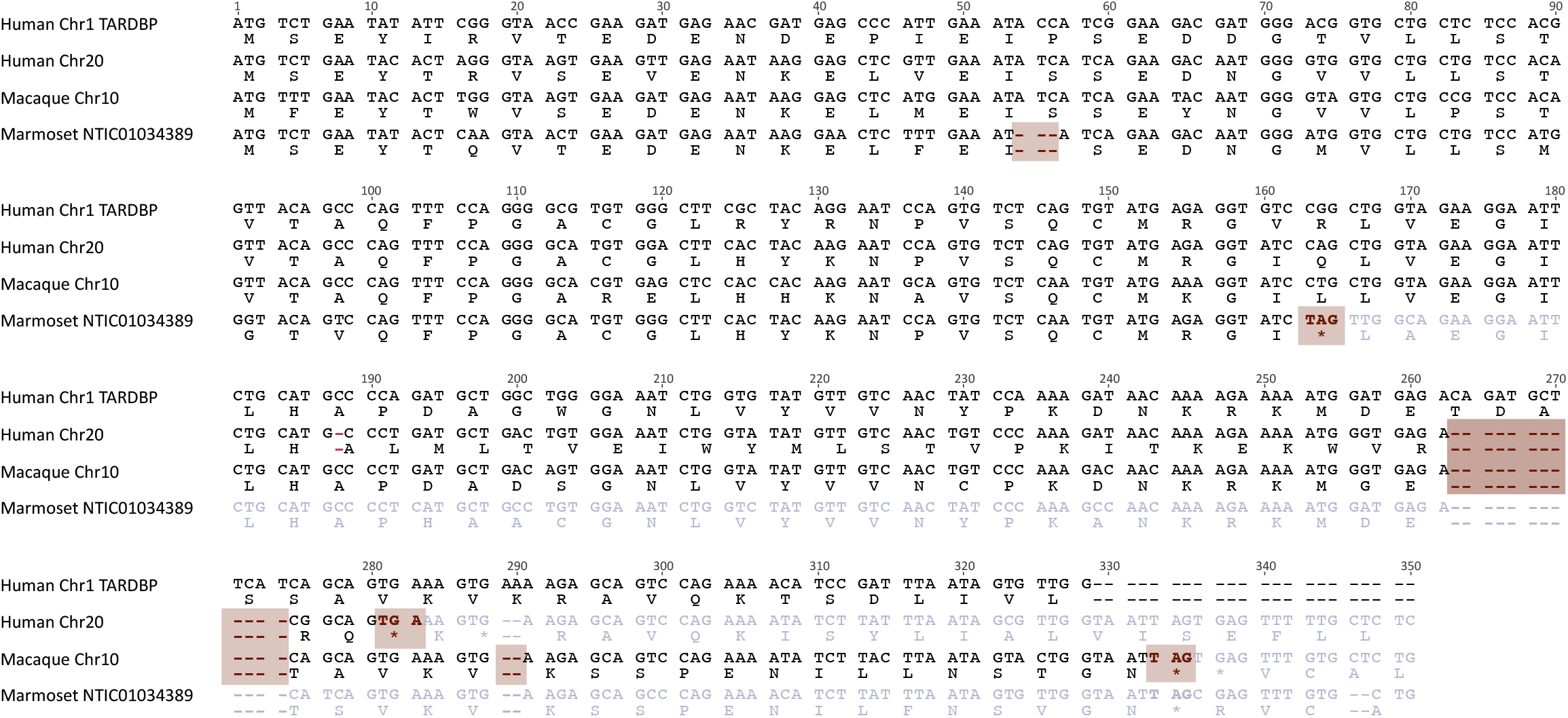
Nucleotide alignment of the TARDBP functional copy of humans (*Homo sapiens*) and TARDBP retrocopies in representative species of primates in which the retrocopy was identified corresponding to the brown lineage in figure 1. The shading highlights the mutations that make the retrocopies non-functional.

**Supplementary figure 5.**
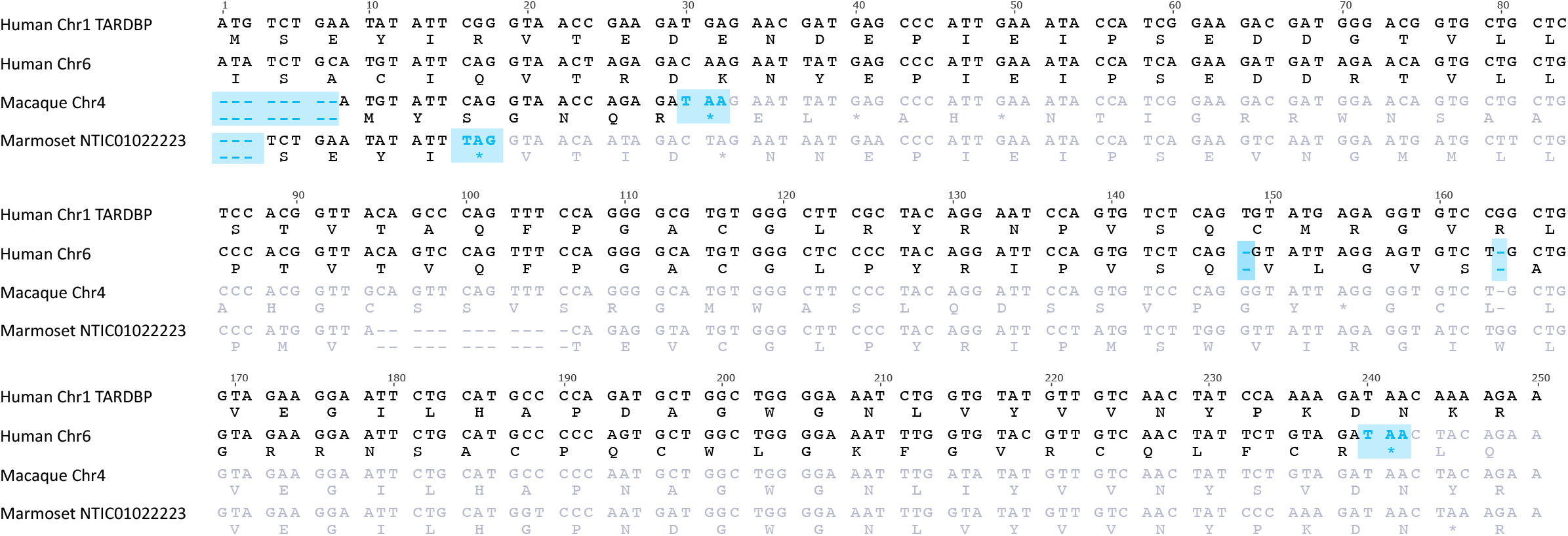
Nucleotide alignment of the TARDBP functional copy of humans (*Homo sapiens*) and TARDBP retrocopies in representative species of primates in which the retrocopy was identified corresponding to the light blue lineage in figure 1. The shading highlights the mutations that make the retrocopies non-functional.

**Supplementary figure 6.**
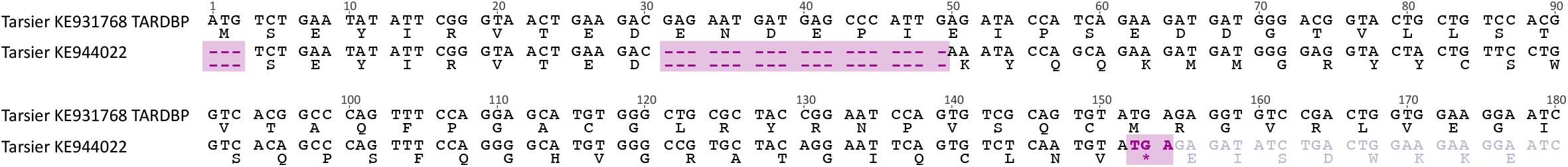
Nucleotide alignment of the TARDBP functional copy of the Philippine tarsier (*Carlito syrichta*) and TARDBP retrocopy identified in the same species. The shading highlights the mutations that make the retrocopy non-functional.

**Supplementary figure 7.**
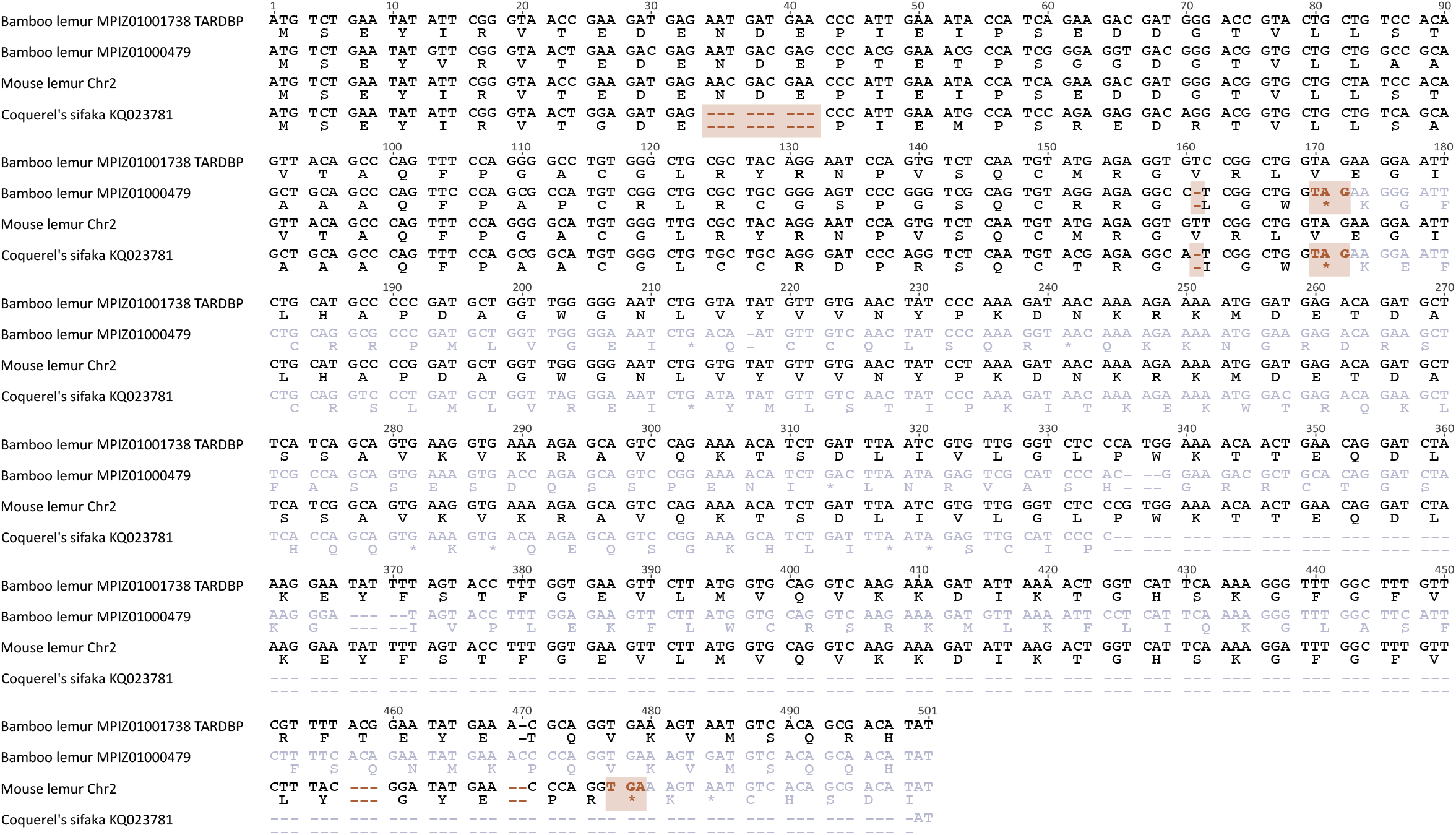
Nucleotide alignment of the TARDBP functional copy of the bamboo lemur (*Prolemur simus*) and TARDBP retrocopies in representative species of strepsirrhines in which the retrocopy was identified on figure 1. The shading highlights the mutations that make the retrocopies non-functional.

## Notes

### Competing Interest Statement

The authors have declared no competing interest.

## References

Altschul SF, Gish W, Miller W, Myers EW, Lipman DJ. 1990. Basic local alignment search tool. J. Mol. Biol. 215:403–410.

Anisimova M, Gil M, Dufayard J-F, Dessimoz C, Gascuel O. 2011. Survey of branch support methods demonstrates accuracy, power, and robustness of fast likelihood-based approximation schemes. Syst. Biol. 60:685–699.

Arai T et al. 2006. TDP-43 is a component of ubiquitin-positive tau-negative inclusions in frontotemporal lobar degeneration and amyotrophic lateral sclerosis. Biochem. Biophys. Res. Commun. 351:602–611.

Casola C, Betrán E. 2017. The Genomic Impact of Gene Retrocopies: What Have We Learned from Comparative Genomics, Population Genomics, and Transcriptomic Analyses? Genome Biol. Evol. 9:1351–1373.

Finstermeier K et al. 2013. A mitogenomic phylogeny of living primates. PLoS One. 8:e69504.

Gearing M, Rebeck GW, Hyman BT, Tigges J, Mirra SS. 1994. Neuropathology and apolipoprotein E profile of aged chimpanzees: implications for Alzheimer disease. Proc. Natl. Acad. Sci. U. S. A. 91:9382–9386.

Gonçalves I, Duret L, Mouchiroud D. 2000. Nature and structure of human genes that generate retropseudogenes. Genome Res. 10:672–678.

Guindon S et al. 2010. New algorithms and methods to estimate maximum-likelihood phylogenies: assessing the performance of PhyML 3.0. Syst. Biol. 59:307–321.

Hanson KA, Kim SH, Tibbetts RS. 2012. RNA-binding proteins in neurodegenerative disease: TDP-43 and beyond. Wiley Interdiscip. Rev. RNA. 3:265–285.

Herrero J et al. 2016. Ensembl comparative genomics resources. Database. 2016. doi: 10.1093/database/bav096.

Heyburn L, Moussa CE-H. 2017. TDP-43 in the spectrum of MND-FTLD pathologies. Mol. Cell. Neurosci. 83:46–54.

Hoang DT, Chernomor O, von Haeseler A, Minh BQ, Vinh LS. 2018. UFBoot2: Improving the Ultrafast Bootstrap Approximation. Mol. Biol. Evol. 35:518–522.

Holt RD, Nesse RM, Williams GC. 1996. Why We Get Sick: The New Science of Darwinian Medicine. Ecology. 77:983. doi: 10.2307/2265522.

Kabashi E et al. 2008. TARDBP mutations in individuals with sporadic and familial amyotrophic lateral sclerosis. Nat. Genet. 40:572–574.

Kaessmann H, Vinckenbosch N, Long M. 2009. RNA-based gene duplication: mechanistic and evolutionary insights. Nat. Rev. Genet. 10:19–31.

Kalyaanamoorthy S, Minh BQ, Wong TKF, von Haeseler A, Jermiin LS. 2017. ModelFinder: fast model selection for accurate phylogenetic estimates. Nat. Methods. 14:587–589.

Katoh K, Standley DM. 2013. MAFFT multiple sequence alignment software version 7: improvements in performance and usability. Mol. Biol. Evol. 30:772–780.

Kay RF, Ross C, Williams BA. 1997. Anthropoid origins. Science. 275:797–804.

Keller MC, Miller G. 2006. Resolving the paradox of common, harmful, heritable mental disorders: which evolutionary genetic models work best? Behav. Brain Sci. 29:385–404; discussion 405-52.

Klim JR, Pintacuda G, Nash LA, Guerra San Juan I, Eggan K. 2021. Connecting TDP-43 Pathology with Neuropathy. Trends Neurosci. doi: 10.1016/j.tins.2021.02.008.

Kumar S, Stecher G, Suleski M, Hedges SB. 2017. TimeTree: A Resource for Timelines, Timetrees, and Divergence Times. Mol. Biol. Evol. 34:1812–1819.

Long M, Betrán E, Thornton K, Wang W. 2003. The origin of new genes: glimpses from the young and old. Nat. Rev. Genet. 4:865–875.

Marques AC, Dupanloup I, Vinckenbosch N, Reymond A, Kaessmann H. 2005. Emergence of young human genes after a burst of retroposition in primates. PLoS Biol. 3:e357.

McDonell L, Drouin G. 2012. The abundance of processed pseudogenes derived from glycolytic genes is correlated with their expression level. Genome. 55:147–151.

Navarro FCP, Galante PAF. 2015. A Genome-Wide Landscape of Retrocopies in Primate Genomes. Genome Biol. Evol. 7:2265–2275.

Neumann M et al. 2006. Ubiquitinated TDP-43 in frontotemporal lobar degeneration and amyotrophic lateral sclerosis. Science. 314:130–133.

Nguyen NTT, Vincens P, Roest Crollius H, Louis A. 2018. Genomicus 2018: karyotype evolutionary trees and on-the-fly synteny computing. Nucleic Acids Res. 46:D816–D822.

Ohshima K et al. 2003. Whole-genome screening indicates a possible burst of formation of processed pseudogenes and Alu repeats by particular L1 subfamilies in ancestral primates. Genome Biol. 4:R74.

Perelman P et al. 2011. A molecular phylogeny of living primates. PLoS Genet. 7:e1001342.

Pinarbasi ES et al. 2018. Active nuclear import and passive nuclear export are the primary determinants of TDP-43 localization. Sci. Rep. 8:7083.

Pozzi L et al. 2014. Primate phylogenetic relationships and divergence dates inferred from complete mitochondrial genomes. Mol. Phylogenet. Evol. 75:165–183.

Robberecht W, Philips T. 2013. The changing scene of amyotrophic lateral sclerosis. Nat. Rev. Neurosci. 14:248–264.

Sreedharan J et al. 2008. TDP-43 mutations in familial and sporadic amyotrophic lateral sclerosis. Science. 319:1668–1672.

Tam OH et al. 2008. Pseudogene-derived small interfering RNAs regulate gene expression in mouse oocytes. Nature. 453:534–538.

Tatusova TA, Madden TL. 1999. BLAST 2 Sequences, a new tool for comparing protein and nucleotide sequences. FEMS Microbiol. Lett. 174:247–250.

Trifinopoulos J, Nguyen L-T, von Haeseler A, Minh BQ. 2016. W-IQ-TREE: a fast online phylogenetic tool for maximum likelihood analysis. Nucleic Acids Res. 44:W232–5.

Uhlén M et al. 2015. Proteomics. Tissue-based map of the human proteome. Science. 347:1260419.

Vamathevan JJ et al. 2008. The role of positive selection in determining the molecular cause of species differences in disease. BMC Evol. Biol. 8:273.

Wang H-Y, Wang I-F, Bose J, Shen C-KJ. 2004. Structural diversity and functional implications of the eukaryotic TDP gene family. Genomics. 83:130–139.

Watanabe T et al. 2008. Endogenous siRNAs from naturally formed dsRNAs regulate transcripts in mouse oocytes. Nature. 453:539–543.

Yates AD et al. 2020. Ensembl 2020. Nucleic Acids Res. 48:D682–D688.

Zhang J. 2003. Evolution by gene duplication: an update. Trends Ecol. Evol. 18:292–298.

Zhang Z, Gerstein M. 2004. Large-scale analysis of pseudogenes in the human genome. Curr. Opin. Genet. Dev. 14:328–335.

Zhao L et al. 2020. TDP-43 facilitates milk lipid secretion by post-transcriptional regulation of Btn1a1 and Xdh. Nat. Commun. 11:341.

